# Cold-acclimation induces life stage-specific responses in the cardiac proteome of Western painted turtles (*Chrysemys picta bellii*): implications for anoxia tolerance

**DOI:** 10.1101/2021.02.03.429598

**Authors:** Sarah L. Alderman, Claire L. Riggs, Oliver Bullingham, Todd E. Gillis, Daniel E. Warren

## Abstract

Western painted turtles *(Chrysemys picta bellii)* are the most anoxia-tolerant tetrapod. Survival time improves at low temperature and during ontogeny, such that adults acclimated to 3°C survive far longer without oxygen than either warm-acclimated adults or cold-acclimated hatchlings. Since protein synthesis is rapidly suppressed to save energy at the onset of anoxia exposure, this study tested the hypothesis that cold-acclimation would evoke preparatory changes in protein expression that would support enhanced anoxia survival in adult but not hatchling turtles. To test this, adult and hatchling turtles were acclimated to either 20°C (warm) or 3°C (cold) for 5 weeks, and then the heart ventricles were collected for quantitative proteomic analysis using labeled isobaric tags and mass spectrometry. The relative abundances of 1316 identified proteins were compared between temperatures and developmental stages. The effect of cold-acclimation on the cardiac proteome was most evident when life stage was included as a covariable, suggesting that ontogenic differences in anoxia tolerance may be predicated on successful maturation of the heart from its hatchling to adult form and, only after this maturation occurs, will cold-acclimation induce protein expression changes appropriate for supporting heart function during prolonged anoxia. The main differences between the hatchling and adult cardiac proteomes reflect an increase in metabolic scope that included more myoglobin and increased investment in both aerobic and anaerobic energy pathways. Mitochondrial structure and function were key targets of life stage- and temperature-induced changes to the cardiac proteome, including reduced complex II proteins in cold-acclimated adults that may help down-regulate the electron transport system and avoid succinate accumulation during anoxia. Therefore, targeted cold-induced changes to the cardiac proteome may be a contributing mechanism for stagespecific anoxia tolerance in turtles.

## Introduction

For most vertebrates, the inability to sustain adequate ATP supply in the absence of oxygen (anoxia) can quickly trigger cell death, and any cells that survive anoxia face the added challenge of mitigating tissue damage related to excess production of reactive oxygen species (ROS) when oxygen returns. These events are particularly devastating to vital organs with high metabolic demands, like the heart. Painted turtles *(Chrysemys picta)* are champions among vertebrates in their capacity to live without oxygen (Jackson, 2002), surviving months at the bottom of ice-covered, anoxic ponds and lakes during their winter dormancy (Jackson and Ultsch, 2010). The ability to survive prolonged periods of anoxia whilst maintaining life-sustaining cardiac function presents a fascinating comparative model for studying the cellular mechanisms of anoxia tolerance and perhaps for identifying novel strategies to preserve cardiac integrity in the event of ischaemia reperfusion.

Painted turtles are well-adapted for anaerobic metabolism and integrate a variety of cellular, physiological and behavioral responses to balance and prioritize energy needs, manage metabolic wastes, and defend vital processes when oxygen is limited, including a profound hypometabolism and bradycardia, large hepatic glycogen stores, and high lactate buffering capacity afforded by the blood and shell (Bickler and Buck, 2007; Hochachka et al., 1996; Jackson, 1968; Jackson, 2000; Jackson and Schmidt – Nielsen, 1966; Overgaard et al., 2007; Stecyk et al., 2008; Storey, 2004). In many tissues, reducing ATP demand is supported to a large extent by a decrease in the energetically expensive process of protein turnover (Bailey and Driedzic, 1996; Land and Hochachka, 1994; Land et al., 1993) facilitated by translational suppression (Fraser et al., 2001; Keenan et al., 2015). In cardiac tissue, however, mechanical work consumes most of the cellular energy, and so reducing heart rate to a minimal level is critical to economizing ATP (Stecyk et al., 2008). Nevertheless, translational arrest does occur in anoxic cardiac tissue (Bailey and Driedzic, 1996), which may be facilitated by a ubiquitous down-regulation of ribosomal protein genes (Fanter et al., 2020). Mitochondrial remodeling also plays a key role in managing metabolic depression (Bundgaard et al., 2020; St-Pierre et al., 2000; Stuart et al., 1998). For example, mitochondrial respiration rates are reduced in the brain and heart of cold-acclimated turtles without altering mitochondrial content or structure (Bundgaard et al., 2019; Pamenter et al., 2016). Rather, suppressing mitochondrial activity may be achieved by a non-specific decrease in substrate utilization across the electron transport system (ETS) (Bundgaard et al., 2019), or via targeted reductions in proton leak and ATP synthase activity (Galli et al., 2013). The capacity for turtles to evoke these responses is influenced by environmental temperature, such that cold-acclimation prior to anoxia exposure substantially decreases the onset of the metabolic insult and increases the survival time (Herbert and Jackson, 1985a; Herbert and Jackson, 1985b; Warren and Jackson, 2007).

Unlike adult painted turtles, hatchlings do not overwinter underwater but rather remain in their terrestrial nests. This overwintering strategy certainly comes with its own challenges (ex. subzero temperatures), but environmental anoxia is not one of them. Hatchlings are able to withstand anoxic bouts, but survival time is far less than in adults (40 d and 170 d at 3°C for hatchling and adults, respectively; (Odegard et al., 2018; Reese et al., 2004)). This contrasts the mammalian paradigm of an age-related decline in anoxia tolerance (Adolph, 1969). Although a smaller mass of mineralized tissue indicates a lower whole-body buffering capacity (Reese et al., 2004), a higher rate of lactate accumulation in the blood of hatchlings also indicates a reduced capacity to suppress metabolism. Recent work highlights that the transcriptional responses of the heart to anoxia exposure (Fanter et al., 2020) and cold acclimation (Fanter et al. in prep) are life stage-specific, suggesting that ontogenic differences, as well as cold-acclimation, may account for the profound anoxia tolerance of adult painted turtles. Few studies have incorporated life stage comparisons into their experimental designs.

To the best of our knowledge, only three studies have endeavored to use proteomics as a tool for understanding the cellular mechanisms of anoxia tolerance in comparative models. Anoxia-induced changes to protein expression were explored in the brains of painted turtles (Smith et al., 2015) and crucian carp *(Carassius carassius;* (Smith et al., 2009)), another anoxia tolerant model species, using 2D-DIGE. Gomez and Richards (2018) determined proteome differences in anoxia-exposed isolated heart mitochondria from *T. scripta,* using a mass spectrometric approach (Gomez and Richards, 2018). Missing from these studies is the contribution of cold-acclimation to the anoxia-tolerant proteome – a notable limitation given that protein synthesis is rapidly suppressed upon anoxia exposure to facilitate metabolic depression (Bailey and Driedzic, 1996; Fraser et al., 2001; Land and Hochachka, 1994; Land et al., 1993), and thus any protein abundance changes necessary for surviving prolonged periods without oxygen should occur prior to the onset of anoxia. Therefore, we tested the hypothesis that cold-acclimation prepares the heart for anoxia exposure by altering cardiac protein expression of adult painted turtles to generate an anoxia-tolerant cardiac phenotype, and that these preparatory changes are blunted or absent in hatchlings. Heart tissue was chosen because cardiac function is necessarily maintained in both life stages irrespective of overwintering strategy and oxygen availability.

## Results

### Characterization of the turtle cardiac proteome

A total of 1844 and 1865 non-redundant protein identifications were generated from each the two iTRAQ 8-plexes, respectively, with 1503 proteins common to both plexes. An additional 182 proteins were removed due to identification by a single unique peptide, and 5 proteins were removed that displayed oppositional abundance values for the internal control sample quantified on both plexes. Therefore, the cardiac proteome of the Western painted turtle was described by 1316 high-quality, non-redundant proteins with abundance values covering 8 orders of magnitude (Supplemental Data S2). The top 50 most abundant proteins in the cardiac proteome, irrespective of life stage or acclimation temperature, included proteins involved in energy metabolism (ex. ATP synthases, creatine kinase), striated muscle contraction (ex. myosin-binding protein C, troponin C), structural integrity (ex. desmin, spectrin alpha chain), as well as storage and transport (ex. myoglobin, apolipoprotein B-100).

Steady state protein expression in the turtle heart was influenced by one or more experimental treatments (absolute fold-change > 1.2, raw p-value < 0.05). In general, protein expression was influenced more strongly by developmental *stage* (Figure 1 A) than by *temperature* (Figure 1B), and this was also true for *interaction* effects, whereby the age response within each acclimation temperature *(stage×temperature;* Figure 1C and 1E) was substantially greater than a temperature response within either age class *(temperature×stage;* Figure 1D and 1F).

**Figure 1.**
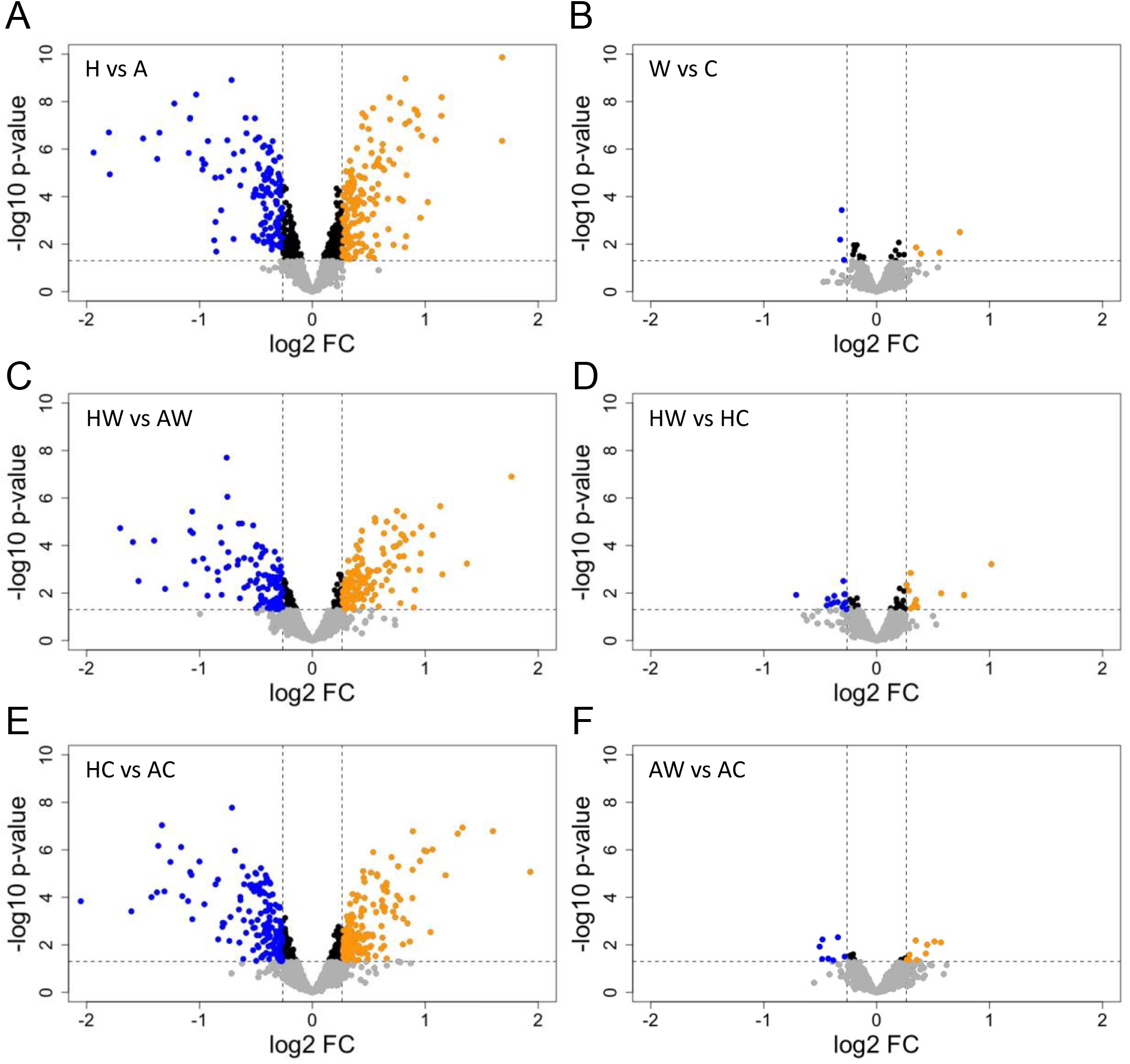
Volcano plots of protein abundance changes across experimental conditions, expressed as the inverse log of the p-value versus log2 fold change (FC). Significant effects were considered for any protein with absolute FC > 1.2 and p<0.05 and are indicated in blue (down-regulated) or yellow (up-regulated). Comparisons used in the analysis included main effects of (A) *age* and (B) *temperature*, as well as interaction effects (*stage*×*temperature; temperature×stage*) including (C) *stage* effects in warm acclimated turtles, (D) *temperature* effects in hatchlings, (E) *stage* effects in cold acclimated turtles, and (F) *temperature* effects in adult turtles. A, adult; C, cold; H, hatchling; W, warm.

### Effect of acclimation temperature on the cardiac proteome

Cold-acclimation had a negligible effect on steady state protein expression in the heart, with just 7 differentially abundant (DA) proteins (0.5% of the quantified cardiac proteome). Furthermore, only gelsolin isoform X1 (log2FC = −0.289) was not influenced by a confounding effect of developmental *stage*. In addition, filamin-C was significantly down-regulated by cold-acclimation, but the response was greater in the hearts of adult than in hatchling turtles (log2FC = −0.343 and −0.278, respectively). Beyond these two proteins, the response of the cardiac proteome to cold-acclimation was stage specific (Table 1), and in the case of carbonic anhydrase 1, oppositional (adult log2FC = 0.512; hatchling log2FC = −0.379). In adult hearts, 9 proteins increased with cold-acclimation, including natriuretic peptides A-like isoform X2, structural proteins (collagen alpha-1(IV) chain isoform X1, integrin alpha-D-like isoform X2), and immune proteins (complement factor H isoform X1, beta-microseminoprotein-like isoform X1), while 7 proteins decreased with cold-acclimation, including contractile proteins (myosin regulatory light chain 10, myosin light chain 3). In contrast, the hatchling-specific response to cold-acclimation included 11 up-regulated proteins such as the serine protease inhibitor, ovomucoid-like isoform X1, and muscle-specific proteins (myosin regulatory light chain 2 atrial isoform, four and a half LIM domains protein 1 isoform X1), while 14 proteins decreased including cell cycle proteins (transforming protein RhoA-like, dynactin subunit 2) and mitochondrial proteins (mitochondrial 2-oxoglutarate/malate carrier protein, NADH dehydrogenase [ubiquinone] 1 alpha subcomplex subunit 5). Functional analysis of these DA proteins did not support the enrichment of cellular pathways based on known protein interactions (no significant PPI networks, GO terms, or KEGG pathways identified in STRING).

**Table 1.** Age-specific responses to cold acclimation. Proteins are listed alphabetically within functional groupings. Significant *temperature×stage* effects are indicated with **bold\*** font for any protein with an absolute log2 fold change (FC) greater than 0.263 and a p-value <0.05. Positive log2FC indicates an increase in protein abundance following cold acclimation in adults or hatchlings, whereas negative values indicate decreased protein abundance.

| Protein | Accession | log2FC |  |
| --- | --- | --- | --- |
|  |  | AW_vs_AC | HW_vs_HC |
| Muscle structure & function |  |  |  |
| calponin-1 | XP_008161545.2 | 0.448* | 0.010 |
| filamin-C | XP_023966385.1 | -0.343* | -0.278* |
| four and a half LIM domains protein 1 isoform X1 | XP_005301111.1 | 0.106 | 0.349* |
| myosin light chain 3 | XP_008168920.1 | -0.428* | 0.227 |
| myosin regulatory light chain 10 | XP_005284475.1 | -0.483* | 0.134 |
| myosin regulatory light chain 2 atrial isoform | XP_005312430.1 | -0.189 | 0.339* |
| myosin-15 isoform X1 | XP_008163413.1 | -0.505* | -0.166 |
| Energy metabolism |  |  |  |
| adenine phosphoribosyltransferase | XP_005292281.1 | -0.011 | 0.285* |
| adenylosuccinate lyase | XP_005302604.1 | 0.115 | -0.268* |
| carbonyl reductase [NADPH] 1 | XP_005303378.1 | 0.292* | 0.032 |
| mitochondrial 2-oxoglutarate/malate carrier protein | XP_005295334.1 | -0.048 | -0.287* |
| NADH dehydrogenase [ubiquinone] 1 alpha subcomplex subunit 5 | XP_005305757.1 | 0.159 | -0.287* |
| phosphomevalonate kinase | XP_005280701.1 | 0.114 | -0.404* |
| Transport |  |  |  |
| band 3 anion transport protein | XP_005302015.1 | 0.049 | -0.433* |
| coatamer subunit beta | XP_008164064.1 | -0.111 | -0.295* |
| coatamer subunit zeta-1 | XP_005307757.1 | 0.345* | -0.001 |
| cysteine-rich protein 1 | XP_005279165.1 | 0.288 | 0.360* |
| hemoglobin subunit pi | XP_023956406.1 | 0.162 | -0.712* |
| importin subunit beta-1 | XP_005301918.1 | 0.082 | -0.375* |
| Structure |  |  |  |
| collagen alpha-1(IV) chain isoform X1 | XP_008164968.1 | 0.267* | 0.152 |
| extracellular matrix protein 1 | XP_005293737.1 | -0.176 | 0.301* |
| integrin alpha-D-like isoform X2 | XP_023960058.1 | 0.355* | -0.204 |
| keratin type II cytoskeletal cochlear | XP_005291931.1 | -0.384* | 0.060 |
| myelin P2 protein | XP_005301574.1 | -0.176 | 0.323* |
| Immune & defence |  |  |  |
| beta-microseminoprotein-like isoform X1 | XP_005311423.2 | 0.286* | -0.047 |
| complement factor H isoform X1 | XP_023957795.1 | 0.435* | 0.246 |
| ficolin-2-like | XP_005300224.1 | 0.230 | 0.571* |
| macrophage migration inhibitory factor | XP_005288622.1 | -0.037 | 0.265* |
| ribonuclease-like | XP_023966987.1 | 0.367 | 0.775* |
| Other |  |  |  |
| amine oxidase [flavin-containing] A | XP_005301830.1 | -0.283* | -0.031 |
| bcl-2-like protein 13 isoform X1 | XP_005288075.1 | -0.480* | -0.188 |
| carbonic anhydrase 1 | XP_005299444.1 | 0.512* | -0.379* |
| carbonic anhydrase 2 | XP_005299442.2 | 0.038 | -0.442* |
| dynactin subunit 2 | XP_008161244.1 | -0.041 | -0.273* |
| elongation factor 2-like | XP_008164272.1 | 0.086 | 0.302* |
| glutathione S-transferase-like | XP_008175413.1 | 0.162 | -0.303* |
| natriuretic peptides A-like isoform X2 | XP_005292869.1 | 0.569* | -0.096 |
| ovomucoid-like isoform X1 | XP_023955986.1 | 0.487 | 1.016* |
| transforming protein RhoA-like | XP_005312008.1 | -0.072 | -0.344* |

### Effect of developmental stage on the cardiac proteome

The cardiac proteome of painted turtles changed substantially as the animals mature from hatchlings to adults. A total of 285 proteins were significantly altered by the main effect *stage,* including calponin-1, fish-egg lectin-like, and myoglobin as the top three up-regulated proteins (highest fold-change; Figure 2A), and fatty acid-binding protein adipocyte, myosin regulatory light chain 2 atrial isoform, and transthyretin as the top three down-regulated proteins (Figure 2B). Most DA proteins for the main effect of developmental *stage,* however, were also significant for a *stage×temperature* interaction effect (Supplemental Data S3), for a combined response of 390 DA proteins. The influence of temperature on the maturation of the cardiac proteome was more substantial in cold-acclimated turtles. Specifically, while ~20% of the total *stage×temperature* DA proteins were unique to the HW_vs_AW comparison, the effect of developmental *stage* in the cold-acclimated turtles claimed up to 40% of the total DA proteins (Figure 3 left panel). This included decreased abundances of three 40S ribosomal proteins (RPS6, RPS21, RPS28) in cold-acclimated adults relative to hatchlings, and increased abundances of two 60S ribosomal proteins (RPL10A, RPL27) in cold-acclimated hatchlings relative to adults (Figure 4).

**Figure 2.**
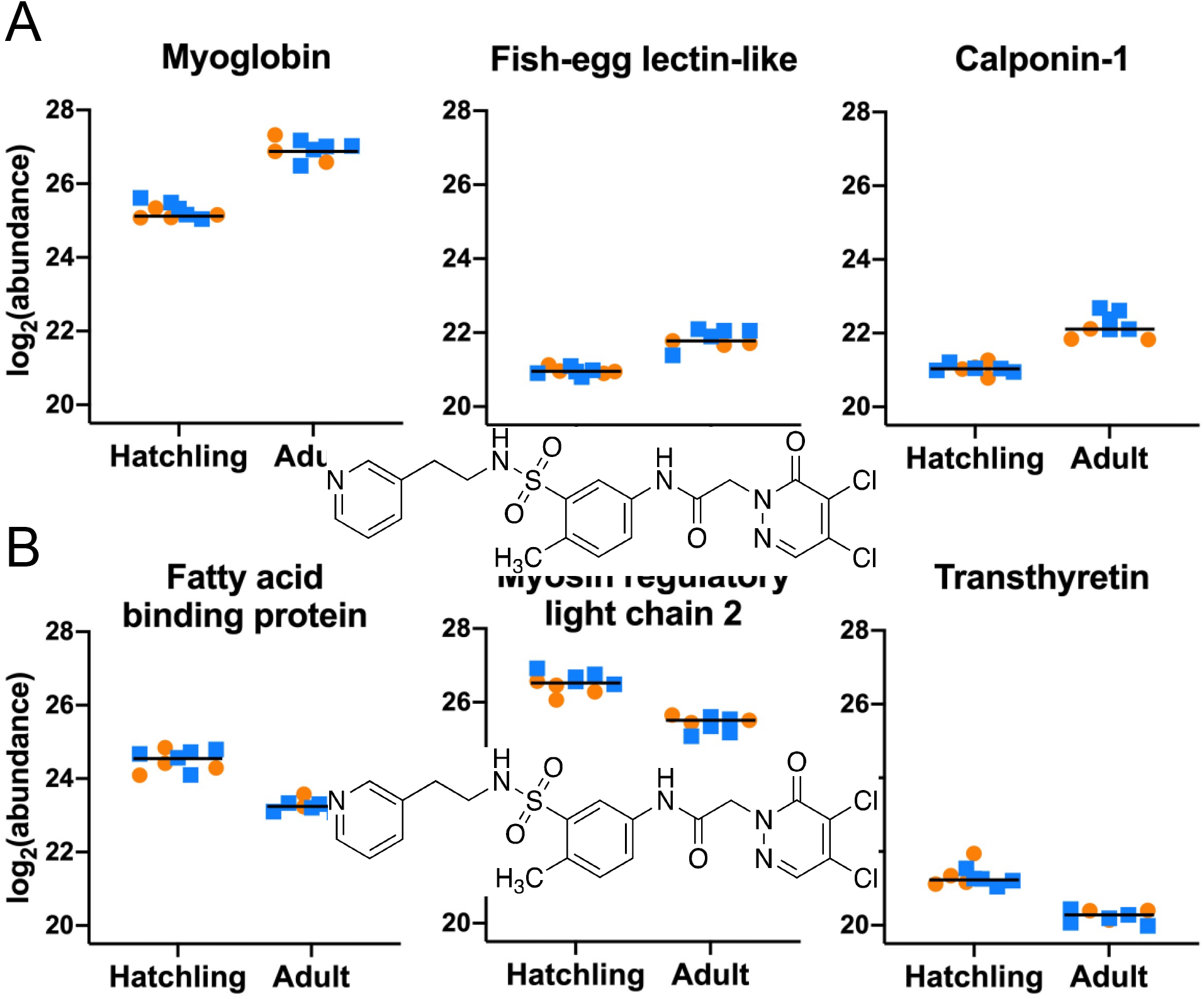
Summary of top differentially abundant proteins for the main effect of *stage* (hatchling vs adult). (A) Top 3 up-regulated proteins. (B) Top 3 down-regulated proteins. Data for individual turtles is plotted as log2(abundance), acclimation temperature is indicated as orange circles (warm) or blue squares (cold), and the line is the geometric mean for *stage* (n=7-8).

**Figure 3.**
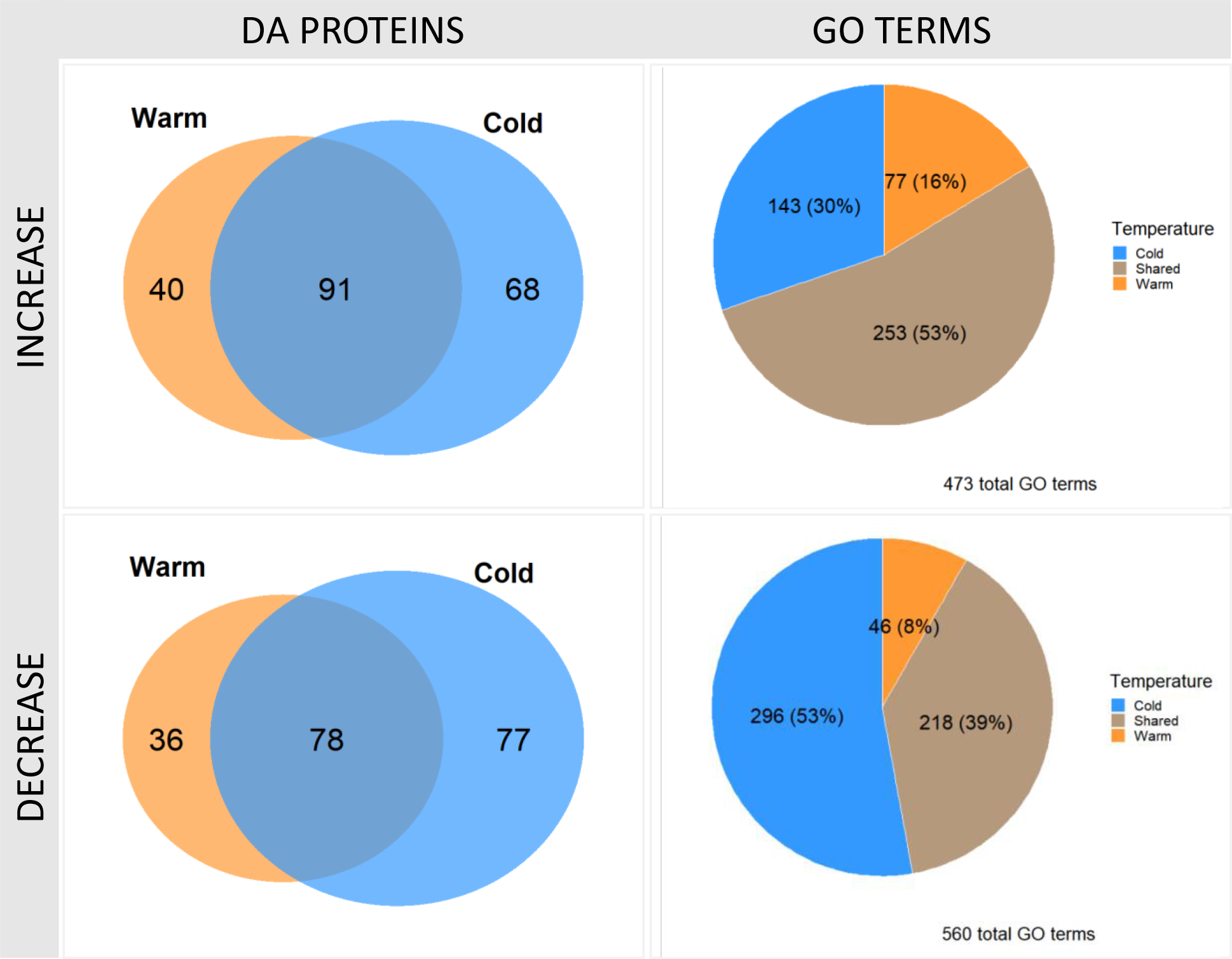
Proportions of shared and unique differentially abundant (DA) proteins (left) and gene ontology (GO) terms (right) for *stage×temperature* interactions in the cardiac proteomes of warm-acclimated hatchling vs warm-acclimated adult (orange), and cold-acclimated hatchling vs cold-acclimated adult (blue). Significant effects are separated for those that increase with age (top) and decrease with age (bottom).

**Figure 4.**
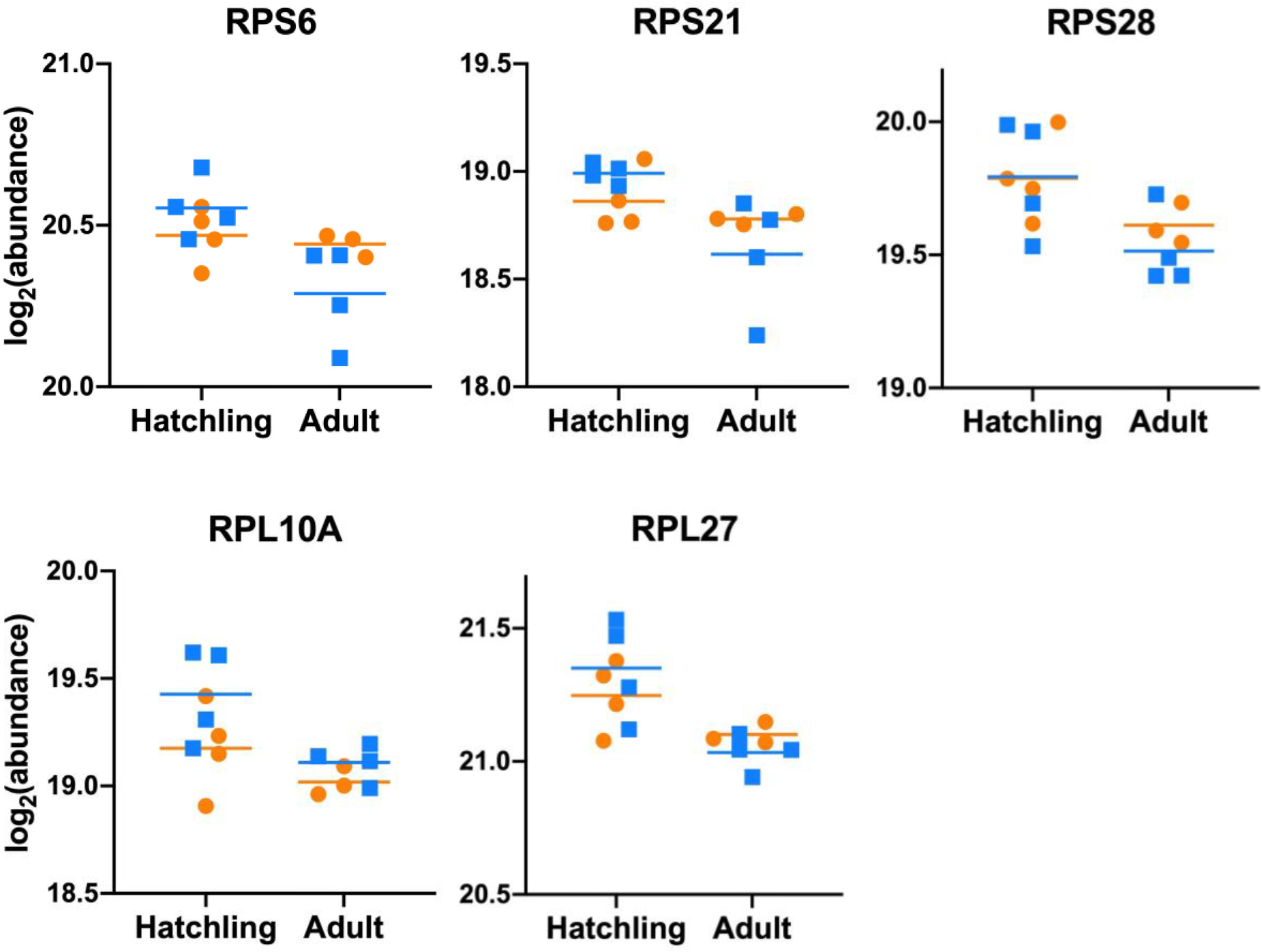
Differentially abundant ribosomal proteins with a significant *stage×temperature* interaction for cold-acclimated turtles. Three 40S ribosomal proteins (RPS6, RPS21, RPS28) had lower abundance in cold-acclimated adult turtle hearts relative to hatchlings. Two 60S ribosomal proteins (RPL10A, RPL27) had greater abundances in cold-acclimated hatchling turtle hearts relative to adults. For each protein, data is plotted as log2(abundance) for warm-acclimated (orange) and cold-acclimated (blue) turtles at each life stage, with geometric mean marked with a coloured line for each *stage* x *temperature* variable. There were no significant differences in the abundances of any of these five proteins for warm-acclimated hatchlings and adults.

Functional insight into the effect of developmental stage on the cardiac proteome was considered separately for each acclimation temperature by inputting the DA proteins for each *stage×temperature* interaction term separately in STRING, and the results interpreted based on shared and unique GO term enrichment. Over 1000 GO terms were enriched by the *stage×temperature* DA proteins (Supplemental Data S4), with a greater proportion of GO term enrichment occurring in the HC_vs_AC, particularly for down-regulated DA proteins. As indicated in Figure 3 (right panel), 39-53% of enriched GO terms were shared between warm- and cold-acclimated turtles, and the majority of the remaining enriched GO terms were unique to cold-acclimated turtles (30-53%). Visualizations of shared and unique GO terms for *stage×temperature* revealed that maturation of the cardiac proteome from hatchling to adults involved a substantial investment in proteins related to mitochondrial form and function (Figure 5). In warm-acclimated adult turtle hearts, enrichment of additional mitochondria-related GO terms occurred relative to warm-acclimated hatchlings, including Cellular Components for the succinate dehydrogenase (SDH) and ATP synthase complexes (Complex II and V, respectively), Molecular Functions associated with oxidoreductase and succinate dehydrogenase activities, and NADH metabolic process (Biological Process; Figure 6 top). In cold-acclimated adult turtle hearts, the majority of GO terms enriched relative to cold hatchlings were categorized as Biological Processes and included up-regulation of pathways involved in purine metabolism and biosynthesis as well as the respirasome (Cellular Component), and down-regulation of pathways involved in RNA processing and biogenesis of cell components (Figure 6 bottom panel). This differential pathway enrichment can be attributed to specific adjustments in ETS Complexes I-IV and ATP synthase (Complex V; Figure 7A). For example, several subunits of Complex I (NADH dehydrogenase [ubiquinone] subcomplex, NDUF) were uniquely altered by *stage* and *temperature* (Figure 7B), including a greater *stage* response in warm (ie. NDUFA6) or cold (ie. NDUFA12, NDUFB9), as well as oppositional responses (ie. NDUFA5). For both Complex II (SDH), Complex III (ubiquinol:cytochrome c reductase, UQCR), and Complex V (ATP synthase), a developmental stage-related increase in certain subunits occurred in warm but not cold acclimated turtle hearts (Figure 7C). Finally, as with complex I, subunits of complex IV (cytochrome c oxidase) had a greater *stage* response in the warm-acclimated (ie. NDUF4A, COX6) and cold-acclimated (ie. COX5A, COX5B) turtle hearts, however here the difference was largely driven by a temperature-induced change in hatchlings rather than adults (Figure 7D). Additional DA proteins contributing to the most significantly enriched GO terms in the cold *stage×temperature* comparison included enzymes involved in purine metabolism (ie. GTP:AMP phosphotransferase, purine nucleoside phosphorylase) and energetic pathways (ie. glyceraldehyde 3-phosphate dehydrogenase, phosphoglucomutase, ATP-citrate synthase) (Supplemental Data S4).

**Figure 5.**
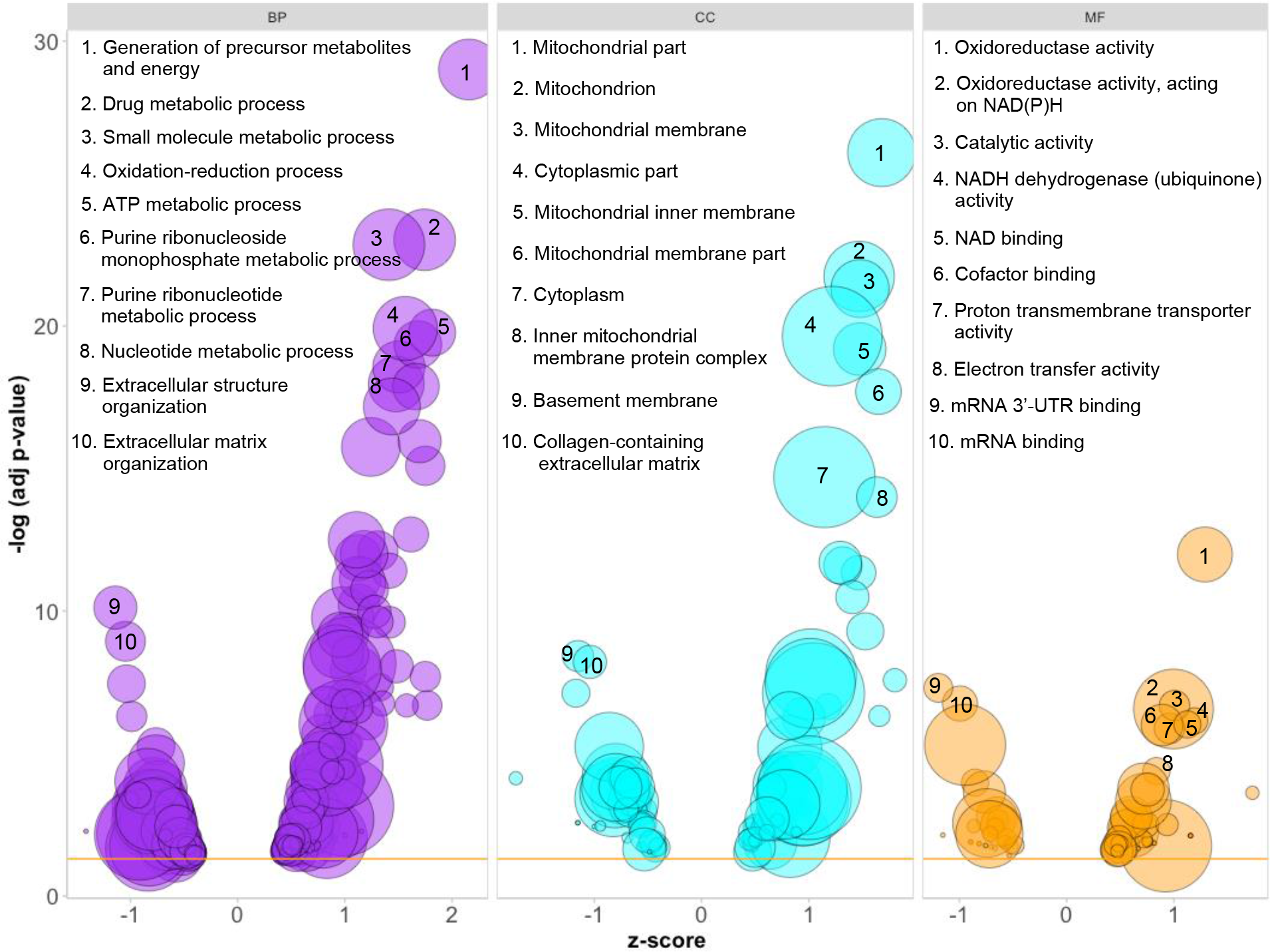
Enriched gene ontology (GO) terms for the main effect of life *stage* and irrespective or acclimation temperature. Data is plotted as the inverse log of the p-value versus z-score. Positive and negative z-scores indicate GO term enrichment for proteins that increase or decrease in abundance from hatchling to adult, respectively. Bubble size corresponds to the number of DA proteins represented in the GO term. The top 8 up-regulated and top 2 down-regulated GO terms are listed for Biological Process (BP; purple), Cellular Component (CC; blue), and Molecular Function (MF; orange).

**Figure 6.**
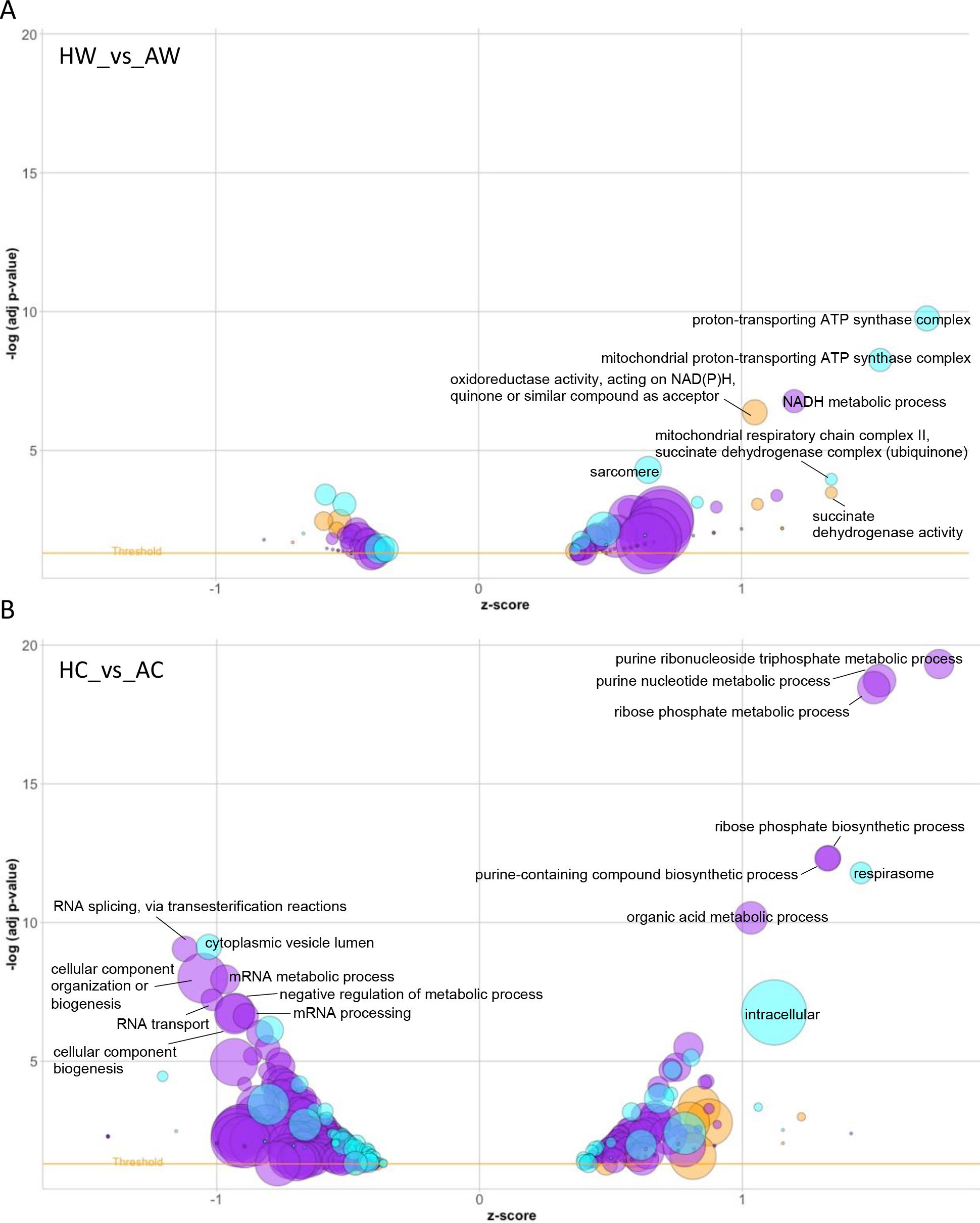
Enriched gene ontology (GO) terms for *stage×temperature* that are unique from a main effect of life *stage*, in (A) warm-acclimated hatchlings relative to adults (HW_vs_AW), and in (B) cold-acclimated hatchlings relative to adults (HC_vs_AC). Data is plotted as the inverse log of the p-value versus z-score. Positive and negative z-scores indicate GO term enrichment for proteins that increase or decrease in abundance from hatchling to adult, respectively. Bubble size corresponds to the number of DA proteins represented in the GO term. Bubble colour corresponds to the primary GO terms Biological Process (purple), Cellular Component (blue), and Molecular Function (orange). A selection of the top up-regulated and down-regulated GO terms are labeled.

**Figure 7.**
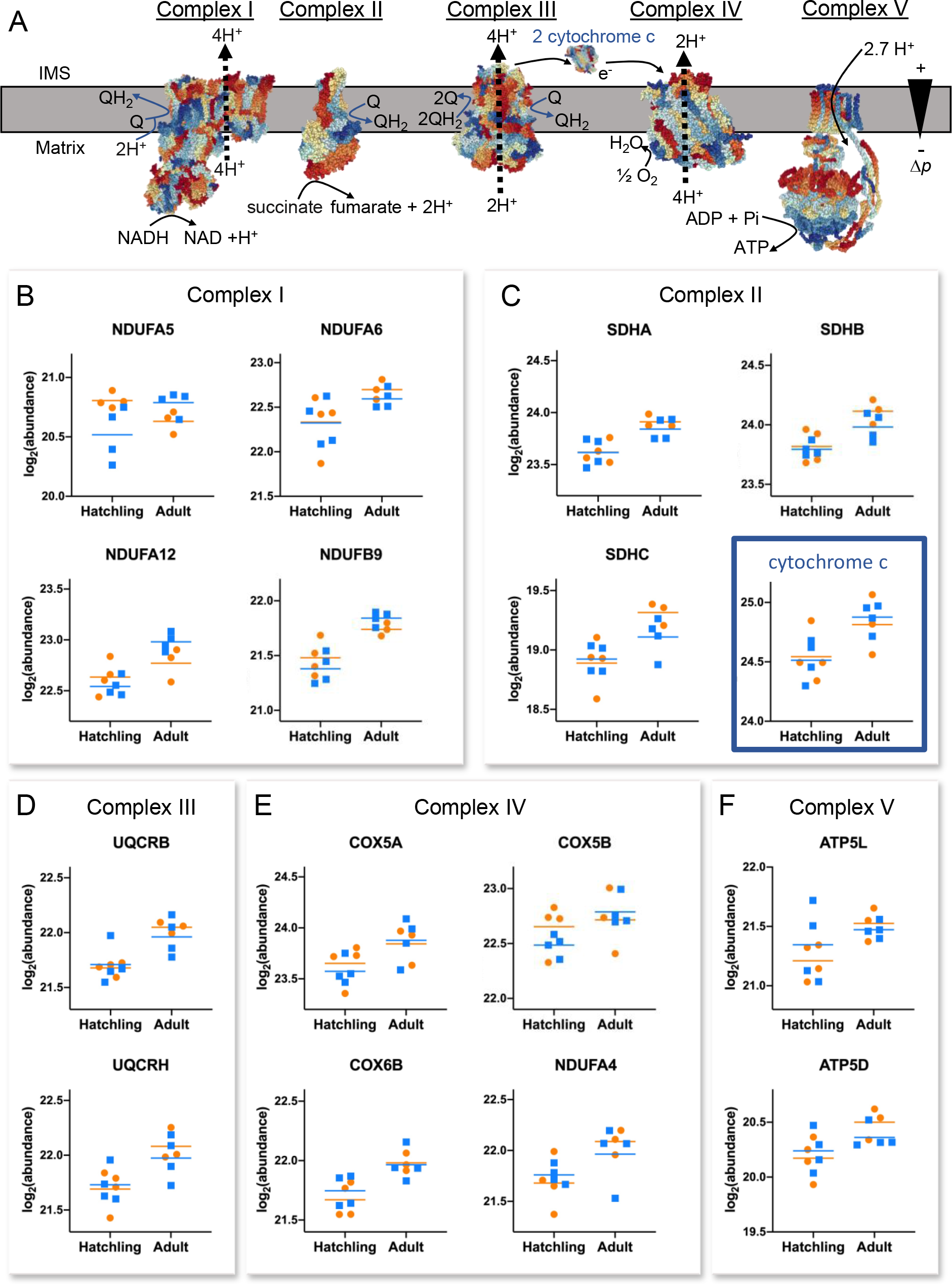
Differentially abundant proteins for *stage×temperature* interaction that are components or accessories of the electron transport system. (A) Schematic of the respiratory chain is modified from (Letts and Sazanov, 2017) using structures from the RCSB Protein Data Bank (complex I (Fiedorczuk et al., 2016), complex II (Sun et al., 2005), complex III (Gao et al., 2003), complex IV (Yano et al., 2016), complex V (Zhou et al., 2015), and cytochrome *c* (Shimada et al., 2017)). **Δ**p indicates the proton motive force, and the intermembrane space (IMS) is oriented down. Complex-specific protein abundances are plotted for individual turtles as log2(abundance) for warm-acclimated (orange) and cold-acclimated (blue) hatchlings and adults, with geometric mean marked with a coloured line for each *stage* x *temperature* variable. Data is grouped by component proteins for complex I (B), complex II (C), complex III (D), complex IV (E), and complex V (F), and cytochrome *c* is outlined in blue. ATP5D, ATP synthase subunit delta; ATP5L, ATP synthase subunit g; COX5A and COX5B, cytochrome c oxidase subunit 5A and 5B; COX6B, cytochrome c oxidase subunit 6B1; NDUFA4, cytochrome c oxidase subunit NDUF4; NDUFA5, NADH dehydrogenase [ubiquinone] 1 alpha subcomplex subunit 5; NDUFA6, NADH dehydrogenase [ubiquinone] 1 alpha subcomplex subunit 6; NDUFA12, NADH dehydrogenase [ubiquinone] 1 alpha subcomplex subunit 12; NDUFB9, NADH dehydrogenase [ubiquinone] 1 beta subcomplex subunit 9; Q, ubiquinone; QH2, ubiquinol; SDHA, succinate dehydrogenase [ubiquinone] flavoprotein subunit; SDHB, succinate dehydrogenase [ubiquinone] iron-sulfur subunit; SDHC, succinate dehydrogenase cytochrome b560 subunit; UQCRB and UQCRH, cytochrome b-c1 complex subunit 7 and 6.

## Discussion

This study provides the first comprehensive and quantitative analysis of protein expression changes incurred by cold-acclimation in the turtle heart, and offers novel, mechanistic hypotheses to explain the developmental stage-specific anoxia tolerance of these animals. A major finding of this work is that maturation of the turtle heart has a more substantial influence on cardiac protein expression than environmental temperature, which has several implications. First, the majority of generalized responses to cold-acclimation so far described for the hearts of painted turtles must result from post-translational modifications rather than changes in the abundance of most proteins themselves. Second, given the pivotal role of the heart in circulating metabolic fuels and wastes, it is perhaps not surprising that painted turtles prioritize the growth and development of the heart even under environmental conditions that may impede heart function. Finally, given the stark differences in overwintering strategies between hatchling and adult painted turtles, and knowing that heart function is necessarily maintained in both scenarios, the ontogenic changes in the cardiac proteome may be a determining factor in preventing hatchlings from spending the winter months submerged in anoxic water.

### Ontogenic changes to the cardiac proteome

There was an overwhelming effect of developmental *stage* on steady-state protein expression in the turtle heart, with as much as 30% of the total identified proteins changing in relative proportion between hatchling and adult hearts. Broadly, these changes involved a downregulation of proteins supporting high-level GO terms related to cell organization and mRNA processing, and up-regulation of proteins associated with energy metabolism, particularly with respect to mitochondrial form and function. These findings are in-line with recent characterizations of the cardiac proteome of American alligators (*Alligator mississippiensis*), where steady-state protein expression in the heart ventricles of late-stage embryos indicated substantial investment in protein synthesis and cellular organization that was not evident in two-year old juveniles (Alderman et al., 2019; Alderman et al., 2020). Combined with the present study, these results emphasize that development of a complex organism is limited by the circulatory system. Thus, protecting a rigid cardiac developmental program ensures that the growth and organization of the heart muscle parallels somatic growth such that increasing peripheral demands from the circulatory system are fulfilled by the capacity of the pump driving the system.

Transthyretin (TTR) was among the most down-regulated proteins associated with maturation in the present study, as was also the case in alligators (Alderman et al., 2020). TTR is a retinol and thyroid hormone carrier protein that is synthesized and secreted by the liver and choroid plexus, and so its repeated detection in the reptilian cardiac proteome is curious. A plausible explanation is that the TTR originated from blood trapped within the spongy ventricular myocardium, as other blood proteins (ex. hemoglobin subunits) were also identified. While the observed age-related decline in TTR abundance is likely not relevant to cardiac maturation, it is consistent with a pronounced ontogenic decrease in circulating TTR across a variety of vertebrate taxa and underscores the importance of thyroid hormones in early life stages (Richardson et al., 2005).

Maturation of the turtle heart included notable investment in proteins involved in muscle energetics, including a concerted investment in ETS component proteins, which would provide an increased capacity for oxidative phosphorylation in adult hearts. Interestingly, elements of the sarcomere were not, in general, proportionally different between life stages, suggesting that the relative number of contractile units does not increase with age in these animals. Therefore, the increased capacity for ATP synthesis as the heart matures would support and sustain faster crossbridge cycling. Myoglobin, a heme protein synthesized by muscle cells, also increased with age in the turtle heart. This would allow the heart to sustain aerobic respiration longer when breath-holding by maintaining a steep oxygen diffusion gradient into the cardiomyocyte and providing a greater capacity for cellular oxygen storage. Oxygen-independent energy pathways were also up-regulated with age in the turtle heart. For example, a life stage-related increase in lactate dehydrogenase A (LDHA) would help defend glycolytic flux rates under oxygen-limiting conditions. Other glycolytic enzymes (ex. fructose biphosphate aldolase C, ALDOC; hexokinase 1, HK1) and creatine kinase M-type (CKM) were also elevated in adult relative to hatchling ventricles. Given that glycolysis is critical for ATP supply during anoxia, it is surprising that the brains of crucian carp and painted turtles reduced one or more of LDH, ALDOC, HK1, and CKM within 24 h of anoxia (Smith et al., 2009; Smith et al., 2015). In carp, LDH protein levels begin to recover towards normoxic levels after 7 d of continuous anoxia (Smith et al., 2009), and so the initial drop in the abundances of glycolytic enzymes may be a transient effect of non-targeted reduction in protein synthesis. Combined, these results suggest that maturation of the turtle heart effectively augments the metabolic scope of cardiomyocytes to support continued cardiac function through daily and seasonal activities. Reciprocally, reduced myoglobin and capacity for aerobic ATP synthesis in hatchling cardiomyocytes, along with a lower hematocrit relative to adults (Fanter et al., 2020), would require that hatchlings switch to anaerobic metabolism sooner and could explain the greater accumulation and longer recovery time for plasma lactate in hatchlings relative to adults under experimental anoxia (Fanter et al., 2020).

### Life stage-specific responses to cold acclimation

Cold-acclimation is recognized for improving anoxia tolerance in turtles and other hypoxia-tolerant species (Boutilier, 2001; Hochachka, 1986). At the same time, an ontogenic threshold for anoxia tolerance likely exists, as adults survive much longer in anoxia than hatchling turtles (Reese et al., 2004). Results of the present study highlight changes in cardiac protein expression that are likely to help generate the more anoxia-tolerant phenotype of cold-acclimated adult turtles relative to hatchlings. These changes include reduced ribosomal protein expression, alterations in component proteins of the ETS, and increased abundance of carbonic anhydrase, which would facilitate regulation of translational processes, mitochondrial respiration, intracellular pH, and ROS production.

A key strategy for surviving anoxia is to reduce energy demand by entering a hypometabolic state (Jackson, 1968; Jackson and Schmidt – Nielsen, 1966). In cardiac tissue, the metabolic costs associated with mechanical work claim the majority of the cellular energy budget, thus the large reductions in heart rate and contractility are crucial to sustaining cardiac function during anoxia (Stecyk et al., 2008). Nevertheless, other cost-saving measures, including translational arrest, are likely important for long-term survival. Cellular and mitochondrial protein synthesis (Bailey and Driedzic, 1996) as well as total ventricular RNA content (Bailey and Driedzic, 1996; Keenan et al., 2015) were reduced in adult turtle cardiomyocytes within hours of anoxia exposure, and a sweeping reduction in transcripts for ribosomal proteins occurred following 20 days of anoxia exposure in cold-acclimated adult turtle hearts (Fanter et al., 2020). Conversely, cardiac protein synthesis appeared to continue unperturbed during 20 h of anoxia at 6°C in hatchling re-eared slider turtle hearts (Brooks and Storey, 1993), and transcription of ribosomal protein genes was elevated after 20 days of anoxia exposure in cold-acclimated hatchling painted turtle hearts relative to normoxia controls (Fanter et al., 2020). Results of the present study showed that cold-acclimation reduced the abundances of three 40S ribosomal proteins in adult hearts relative to hatchlings, and two 60S ribosomal proteins were elevated in cold-acclimated hatchling hearts relative to adults. This suggests that the cellular energy savings that come from reducing translational activities may already be at play during cold-acclimation and prior to the onset of anoxia, and that the absence of this response in the hatchling heart may therefore contribute to the lower anoxia tolerance at this life stage.

An interaction between life stage and acclimation temperature was observed for several ETS components, ATP synthase, and cytochrome *c*; however, the responses were asymmetrical and complex-specific. For example, several complex I component proteins (NDUFA5, NDUFA12, NDUFB9) were oppositely affected by cold-acclimation in hatchling and adult turtle hearts. These findings are interesting to consider from the perspective of impending oxygen deprivation in adult but not hatchling turtles. For hatchlings, oxygen supply is unlikely to become limiting in their terrestrial overwintering environment, and so oxidative phosphorylation could continue as the primary supplier of ATP. Therefore, any cold-induced changes to components of the ETS may serve to compensate for the stabilizing effect of reduced temperature on protein structure and support the continued activity of the ETS at low temperature. Notably, hatchlings appear to switch reliance from COX5 to COX6 during cold acclimation, an effect not observed in adults. As cytochrome *c* oxidase (complex IV) employs molecular oxygen as the final electron acceptor in the ETS, a hatchling-specific response to this complex supports its sustained dependency on oxygen during winter. For adult turtles, observed changes in ETS component proteins induced by cold-acclimation may play a role in the metabolic preparations that enhance anoxia survival at low temperature. For example, the adult-specific decrease in several component proteins of complexes II, III, and V with cold-acclimation would reduce the overall aerobic capacity of the heart and may contribute, at least in part, to the observed reduction in respiration rates of isolated mitochondria (Bundgaard et al., 2019; Galli et al., 2013) and the lower cellular ATP content (Stecyk et al., 2009) of cold-acclimated adult turtles.

The switch to anaerobic production of ATP leads to metabolic acidosis from accumulation of lactic acid. At the whole animal level, turtles manage this acidosis by having a comparatively high whole-body buffering capacity, particularly by the shell, which releases buffer and sequesters more than half of the lactate load (Jackson, 2000). In isolated myofilaments from turtle hearts, calcium sensitivity and development of tension were reduced at low pH (Fanter et al., 2017), suggesting that the decrease in intracellular pH (pHi) associated with anoxia may serve as an endogenous mechanism for reducing contractility – and by extension energy requirements – of the heart. Results of the present study suggest that carbonic anhydrase (CA), a key enzyme for regulating pHi, may play a key role given the oppositional effects of cold-acclimation on CA isoform abundance in hatchling and adult turtle hearts. Specifically, CAI and CAIX were up-regulated in the hearts of cold-acclimated adults, while CAI and CAII were down-regulated in cold-acclimated hatchlings. CA catalyzes the reversible hydrolysis of CO_2_ to form HCO_3_^-^ and H^+^, and the fast kinetics of this enzyme, coupled with the diffusibility of CO_2_ across membranes, provides cells with a rapid and efficient means for regulating pHi as well as numerous pH-dependent physiological processes (ex. (Gilmour and Perry, 2009; Maren, 1967; Purkerson and Schwartz, 2007)). Interestingly, mammalian cancer cells rely on membrane-bound CAIX and cytoplasmic CAII to stabilize pHi in glycolytic tumor cells, allowing them to divide and invade into surrounding hypoxic tissue (Pastorekova and Gillies, 2019). At the same time, CAIX also facilitates lactate extrusion from the tumor cell through non-catalytic cooperation with a lactate/H^+^ cotransporter (Pastorekova and Gillies, 2019). The cold-induced increase in CAIX and CAI (cytoplasmic) in adult turtle hearts would establish a similar capacity for managing metabolic acidosis, and therefore may represent preparatory response for impending anoxia that is noticeably absent in hatchlings.

For adult turtles, the temporal variations in temperature and oxygen availability through the winter season mean that not only do cold temperatures precede anoxia, but they also persist as lake surfaces thaw and atmospheric oxygen becomes accessible again. It is interesting, therefore, to consider the cardiac proteome of cold-acclimated adults with respect to how these changes may help to limit tissue damage upon reoxygenation. In mammals, reoxygenation after ischaemia drives a reversal of electron transfer as a consequence of ADP depletion and succinate accumulation during anoxia, leading to overproduction of ROS and triggering apoptosis (Chouchani et al., 2014; Murphy, 2009). Specifically, complex V is unable to dissipate the proton motive force when oxygen returns to the system without an adequate supply of adenine nucleotides, forcing electron transfer backwards from complex II to complex I and generating superoxide (Chouchani et al., 2014; Chouchani et al., 2016). Yet despite months in anoxic conditions, the adult turtle is able to avoid tissue damage upon reoxygenation, suggesting a highly effective ROS management strategy. High constitutive levels of antioxidant enzymes may be a factor in this strategy [49-51, but see 19], and the recently described peroxidase activity of myoglobin (Mannino et al., 2019) coupled with the abundance of this protein in the turtle heart (present study) support a role for myoglobin in antioxidant defense of the turtle heart. In addition to managing ROS damage, the turtle heart also minimizes ROS production upon reoxygenation by limiting succinate accumulation (Buck, 2000; Bundgaard et al., 2018) and stabilizing ADP levels (Bundgaard et al., 2019; Galli et al., 2013) during the anoxic interval. Results of the present study suggest that targeted protein abundance changes during cold-acclimation may be a contributing mechanism in limiting ROS production. In adult but not hatchling turtle hearts, cold-acclimation induced a down-regulation of several complex II component proteins (SDHA, SDHB, SDHC) which would help to slow succinate accumulation in anoxia. At the same time, GO terms related to purine metabolic processes were the top enriched pathways in cold-acclimated adults, suggesting enhanced efforts to regulate adenine nucleotide levels to maintain a balance in intracellular ATP/ADP.

### Conclusions

Physiological investigations of painted turtles and other anoxia-tolerant vertebrates have resulted in a comprehensive and integrative understanding of how animals can survive without oxygen. Onto this long history of research, the present study maps a holistic perspective of protein-level changes in the turtle heart that underpin a rigid developmental program and a temperature-induced phenotype ultimately capable of withstanding months of anoxia. Importantly, results of this study highlight the crucial importance of ontogeny in bestowing anoxia tolerance in turtles, and so future efforts to better define the functional properties of the hatchling turtle heart relative to adults may yield new insights into cardiac function during and following anoxia exposure.

## Materials and Methods

### Animal procurement & husbandry

Adult male Western painted turtles *Chrysemys picta bellii* (N = 7, average mass = 166.2 ± 7.6 g, range = 141.7 – 193.1 g) were purchased from Niles Biological, Inc. (Sacramento, CA, USA) and acclimated for 50 days in large, flow-through fiberglass tanks containing dechlorinated St. Louis municipal water (pH 7-8.5, 13-20°C), with access to deep water and a basking platform bathed in 10 W UV fluorescent light (Zoomed Reptisun 10.0) and 60 W incandescent light maintained on a weekly-updated natural photoperiod. Air temperature was 20-24°C. Adult turtles were fed commercial turtle pellets (Mazuri^®^ Aquatic Turtle Diet) *ad libitum* five times per week.

Painted turtle eggs were collected in June from multiple nest sites located in Rice Creek Chain of Lakes Park (Anoka County, MN; DNR Fisheries Special Permit 23851). Clutches were transferred to containers with moistened grade-3 vermiculite (1.12 g sterile deionized water per g vermiculite) and incubated at 25°C in a field incubator for 3-7 days (to produce all males). Eggs were then incubated at 25°C and ~95% humidity until hatch (8-9 weeks), after which hatchlings were single-housed at 20°C in black plastic containers with a moistened gauze pad until used in experiments in November. All housing and animal procedures were approved by Saint Louis University (Protocol 2198).

### Experimental design and turtle sampling

The experimental set-up, temperature acclimation, and sampling were carried out as previously described (Fanter et al., 2020) in 170 L glass aquarium within which a 10.5 L acrylic aquarium with plastic grating floor and walls was glued to the inside wall. This allowed hatchling and adult turtles to be physically isolated whilst sharing environmental conditions. The tank was partially filled with dechlorinated, aerated, St. Louis municipal water, partially refreshed every 10 days, and no basking platform was provided. A winter photoperiod (9:15 light:dark) was maintained with a 60 W incandescent bulb over the tank. Water temperature was controlled with a proportional temperature controller (YSI model 72) and immersion water heater. The system was housed within a walk-in cold room that was always 1°C colder than set temperature of the YSI controller. Food was withheld because turtles do not eat when their body temperatures are below 15°C.

In November, hatchling turtles (N = 8, average mass = 4.8 ± 0.3 g, range = 3.8 – 5.6 g) were transferred from their individual containers in the laboratory incubator to the acclimation aquarium containing 20°C water. Adult turtles were placed in the outer tank 10 days later. Water level was always maintained to yield a ratio of 51.8 grams of turtle per 1 liter of water. All turtles were then held at 2Ü°C for four more days before sampling a subset as described below (n=3 adult, n=4 hatchling). For the remaining turtles, water temperature was gradually cooled at a rate of 2°C/d to 1Ü°C and then by 1°C/d to 3°C. Turtles were then maintained at 3°C for an additional five weeks prior to sampling. There were no significant differences in body mass between the two temperature groups (2Ü°C and 3°C) for each developmental stage (adults: N = 7, unpaired t-test, P = 0.68; hatchlings: N = 8, unpaired t-test, P = 0.62). To eliminate potential clutch effects, hatchlings were distributed between the two acclimation temperatures (2Ü°C and 3°C) so that all turtles within a treatment group were from unique clutches.

### Tissue Sampling

Turtles were sampled at 20°C and 3°C as previously described (Fanter et al., 2020). At each temperature, the turtles were removed from the water and quickly decapitated, after which the plastrons were removed with scissors (hatchlings) or a bone saw (adults). The heart ventricle was then removed, blotted to remove excess blood, trisected (adults only), and then freeze-clamped in liquid-nitrogen. Ventricles were stored at −80°C until shipping overnight on dry ice to the University of Guelph for proteomics analysis.

### Protein extraction and iTRAQ labelling

A total of 15 frozen ventricles were used to characterize and quantify the cardiac proteome of turtles at two developmental stages (hatchling, H; adult, A) and two acclimation temperatures (warm acclimated, W; cold acclimated, C) for a factorial sample distribution (n=4 each for HW and HC; n=4 AC; n=3 AW). Frozen hearts were homogenized in 150-450 μl icecold RIPA buffer (150 mM NaCl, 50 mM Tris, 100mM DTT, 1% Triton X-100, 0.5% sodium deoxycholate, 0.1% SDS, pH 8) containing a cocktail of protease inhibitors (Promega Corporation, Madison, WI) using a Precellys24 and 2 mm zirconium oxide beads (2 x 25 s at 6800 rpm; Bertin Instruments, Montigny-le-Bretonneux, France). To facilitate dissociation of membrane proteins, the crude homogenate was incubated on ice for 30 min with periodic mixing and a brief sonication (3x 2 s) halfway through the incubation, then clarified by centrifugation (13 000 g x 5 min at 4°C). An aliquot of the supernatant was used to confirm extraction efficacy using standard SDS-PAGE and Coomassie stain. The remaining supernatant was precipitated using the Calbiochem Protein Extraction Kit (EMD Millipore, Billerica, MA) according to manufacturer’s instructions, and reconstituted in 20-50 μl buffer (1 M HEPES, 8 M urea, 2 M thiourea, 4% w/v CHAPS, pH 8.5). For each sample, 150 μg total protein was transferred to an

Amino Ultra-0.5-centrifugation filter device (10K nominal molecular weight cut-off) for buffer exchange, cysteine blocking (8 M urea, 0.1 M HEPES, 0.5 M iodoacetamide, pH 8.5; 20 min in dark), and overnight protein digestion using 4 μg MS-grade trypsin (Promega) at 37°C, all as previously described (Alderman et al., 2019; Dindia et al., 2017). Digested peptides were recovered and labelled using two 8-plex iTRAQ kits (SCIEX, Framingham, MA) according to manufacturer’s instructions. One sample (AW) was labelled in replicate reactions to serve as an internal control on the 2 plexes. Labeled peptides were pooled and purified using C18 columns (Sigma) and eluted with 70% acetonitrile containing 0.1% formic acid.

### Separation of peptides and LC mass spectrometric analysis

Analysis of iTRAQ-labelled peptides by mass spectrometry was performed at SPARC BioCenter Molecular Analysis (The Hospital for Sick Children, Toronto, ON) using an Orbitrap Fusion Lumos (Thermo Scientific), with specifications detailed in Supplemental Data S1. Identification and quantification of cardiac proteins was performed using Proteome Discoverer v2.2.0.388 (Thermo-Fisher, Waltham, MA) by searching mass spectra against the Uniprot non-redundant protein database (February 7, 2020) using the following spectrum file search settings: 20 ppm precursor mass tolerance, 0.5 Da fragment mass tolerance, carbamidomethylation and iTRAQ static modifications, at least 2 unique identifying peptides, and target false discovery rate (FDR) 5%. Protein abundances were normalized to total protein in each run. Data is available at ProteomExchange PXD023534.

### Data processing and bioinformatics

The full list of retrieved protein identifications was collated to remove duplicated entries, proteins not present on both plexes, and proteins identified by single unique peptides. Abundance values for each plex were then normalized using a variance stabilization function (vsn) and missing values imputed using k-nearest neighbour classification (knn). Data from the two plexes were combined and abundance values scaled to the internal control sample. All data processing steps were performed in R Studio. Differentially abundant (DA) proteins were identified using the Differential Enrichment analysis of Proteomics data package in R (DEP 1.10.0), accounting for the influence of the main effects of life *stage* (H, A) and *temperature* (W, C), as well as their *interaction* (*stage×temperature*: HW_vs_AW and HC_vs_AC; *temperature×stage*: HW_vs_HC and AW_vs_AC). To maximize input for functional analyses, significant differences were considered for any protein with an absolute fold-change in abundance greater than 1.2 and a raw p-value less than 0.05 in one or more of the above comparisons. Pathway analysis was performed by uploading the DA protein lists into STRING (https://string-db.org; July 20, 2020) and enriched GO terms were visualized with GOplot (Walter et al., 2015) using a calculated z-score as a measure of term enrichment that accounts for the proportion of the GO term represented by DA proteins in the data set (where z-score = observed / √background, with a negative integer for down-regulated terms).

## Acknowledgments

The authors thank Dr. C. Fanter (Saint Louis University) for assistance conducting the overwintering experiment, and Dr. K. Cottenie (University of Guelph) for help writing R script. S.L.A. and T.E.G. are supported by Natural Sciences and Engineering Research Council (NSERC) of Canada Discovery Grants. O.B. received a Morwick Summer Research Assistantship (University of Guelph) and a Mitacs Undergraduate Research Training Award. This study was funded by a National Science Foundation CAREER grant (award #1253939) to D.E.W.

